# DeepVelo: Deep Learning extends RNA velocity to multi-lineage systems with cell-specific kinetics

**DOI:** 10.1101/2022.04.03.486877

**Authors:** Haotian Cui, Hassaan Maan, Michael D. Taylor, Bo Wang

**Affiliations:** Department of Computer Science, University of Toronto, Toronto, ON, Canada; Vector Institute, Toronto, ON, Canada; Peter Munk Cardiac Centre, University Health Network, Toronto, ON, Canada; Department of Medical Biophysics, University of Toronto, Toronto, ON, Canada; Department of Laboratory Medicine and Pathobiology, University of Toronto, Toronto, ON, Canada; Departments of Surgery, University of Toronto, Toronto, ON, Canada; Division of Neurosurgery, The Hospital for Sick Children, Toronto, ON, Canada; The Arthur and Sonia Labatt Brain Tumor Research Centre, The Hospital for Sick Children, Toronto, ON, Canada

## Abstract

The introduction of RNA velocity in single-cell studies has opened new ways of examining cell differentiation and tissue development. Existing RNA velocity estimation methods rely on strong assumptions of predefined dynamics and cell-agnostic constant transcriptional kinetic rates, which are often violated in complex and heterogeneous single-cell RNA sequencing (scRNA-seq) data. To overcome these limitations, we propose DeepVelo, a novel method that estimates the **cell-specific** dynamics of splicing kinetics using Graph Convolution Networks (GCNs). DeepVelo generalizes RNA velocity to cell populations containing time-dependent kinetics and multiple lineages, which are common in developmental and pathological systems. We applied DeepVelo to disentangle multifaceted kinetics in the processes of dentate gyrus neurogenesis, pancreatic endocrinogenesis, and hindbrain development. The method infers time-varying cellular rates of transcription, splicing and degradation, recovers each cell’s stage in the underlying differentiation process, and detects functionally relevant driver genes regulating these processes. DeepVelo relaxes the constraints of previous techniques, facilitates the study of more complex differentiation and lineage decision events in heterogeneous scRNA-seq data, and is more computationally efficient than previous techniques.

## 2 Main

The concept of RNA velocity refers to the time derivative of the mRNA abundance in a cell, which reflects the changing rate of RNA processing and degradation. Current velocity estimation methods leverage the observation that the abundance and ratio between unspliced pre-messenger RNAs and spliced mature messenger RNAs can be used to infer changes in gene expression dynamics. Higher abundance and ratio of unspliced mRNAs to spliced mRNAs indicates increasing transcription of a certain gene – in other words, up-regulation/induction and a high velocity estimate. Conversely, a lower abundance and indicated ratio lead to a low velocity estimate associated with down-regulation/repression. An equilibrium phase occurs when this dynamical process reaches a stable steadystate. Since unspliced mRNAs can be distinguished in common single-cell RNA sequencing (scRNA-seq) protocols [16], the idea of estimating dynamic RNA velocity using only static sequencing libraries becomes feasible.

The original RNA velocity approach [16] utilized the assumption that the observed transcriptional phases in scRNA-seq last long enough to reach both an apex of induction and a quiescent steady-state equilibrium. This technique infers a per-gene *steady-state ratio* using linear regression, and then RNA velocities are calculated as the deviation of the observed ratio from the steady-state level. This workflow implies two underlying assumptions, (1)**the assumption of steady-state**: For every gene, sufficient number of sequenced cells are at the steady states; (2)**the assumption of cell-agnostic kinetic rates**: The degradation and splicing rate for each gene is shared across all cells. These assumptions are often violated in complex biological systems and bring about limitations in downstream applications, particularly when cell states are partially observed or undergo transcription dynamics more complex than the steadystate pattern. Although a later approach, scVelo [4], attempted to generalize the *steady-state* assumption by replacing these states with *four transcriptional states* and modeling them with a dynamical model, the aforementioned second limitation still remains. Further, scVelo assumes a cyclic trajectory within the four transcriptional states for all observed genes, but this assumption also rarely holds in real-world single-cell datasets with complex differentiation trajectories and multifactorial kinetics [9]. Although several related works have been further developed, including MultiVelo [20], Chromatin Velocity [26], protaccel [8] for extending Velocity beyond RNA, VeloAE [24] for denoising velocity with Deep Neural Nets, Dynamo [25] for exploiting the metabolic labeling sequencing data, the core velocity computation follows the original ideas and therefore the aforementioned limitations still hold.

Overall, existing techniques assume each gene follows a pre-defined trajectory depicted by constant cell-agnostic kinetic rates. This workflow implies that each gene goes through the same velocity trajectory across all celltypes, and limits the application in complex cell systems. To resolve these limitations, we highlight the need for *cell-specific kinetics* which enables the modeling of multilineage systems with heterogeneous cell populations. We propose DeepVelo, a deep neural network based method for RNA velocity estimation. (1) DeepVelo is optimized using a newly introduced probabilistic learning framework, resulting in an approach that is unbiased from pre-defined kinetic patterns. (2) Empowered by Graph Convolutional Networks (GCN), DeepVelo infers **genespecific and cell-specific** RNA splicing and degradation rates. Therefore, compared with the cell-agnostic parameters used in existing techniques [16, 4], DeepVelo is able to model RNA velocity for differentiation dynamics of high complexity, particularly for cell populations with heterogeneous celltypes and multiple lineages.

We demonstrate the efficacy of DeepVelo on multiple developmental scRNA-seq datasets, including dentate gyrus neurogenesis [11], pancreatic endocrinogenesis [2], and hindbrain development [30]. DeepVelo yields more consistent velocity estimates and accurately identifies transcriptional states than existing models. We examine the estimated kinetic rates for individual genes and show that the cell-specific rates accurately recover known differentiation trajectories in challenging scenarios of time-dependent and multi-trajectory gene regulation dynamics. For downstream tasks, DeepVelo can identify putative driver genes of these transcriptional changes, which are more likely to characterize and be involved in dictating lineage fate-decisions. The DeepVelo method is available at (https://github.com/bowang-lab/DeepVelo).

## 3 Results

### 3.1 The DeepVelo model

Modeling the transcriptional dynamics in single cells provides the theoretical basis of RNA velocity. For each cell, the dynamics of transcription, splicing, and degradation (Fig. 1a) can be approximated as the following differential processes

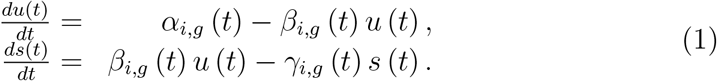

where *α_i,g_, β_i,g_, γ_i,g_* are the kinetic rates for cell *i* and gene *g. t* denotes a time coordinate in cell development. Unspliced immature mRNA is first generated by transcription of DNA and then post-transcriptionally modified and spliced into mature RNA. The dynamics of unspliced RNA abundance, 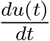, is modeled by the first equation where *α_i,g_* and *β_i,g_* denote the rates of transcription and splicing, respectively. Similarly, the second equation models the dynamics of spliced RNA abundance, 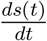, and *γ_i,g_* denotes the rate for RNA degradation. Note these the kinetic rates are intrinsically cell-specific since there is a high degree of variability in transcriptional dynamics between cells [12]. Further, these intrinsic cell-specific transcriptional dynamics are likely to be similar among similar celltypes [19], necessitating celltype-specific parameters. **However, previous velocity estimation techniques** [16, 4] assume global constant kinetic rates across cells, leading to limitations in inferring multi-lineage dynamics.

**Figure 1:**
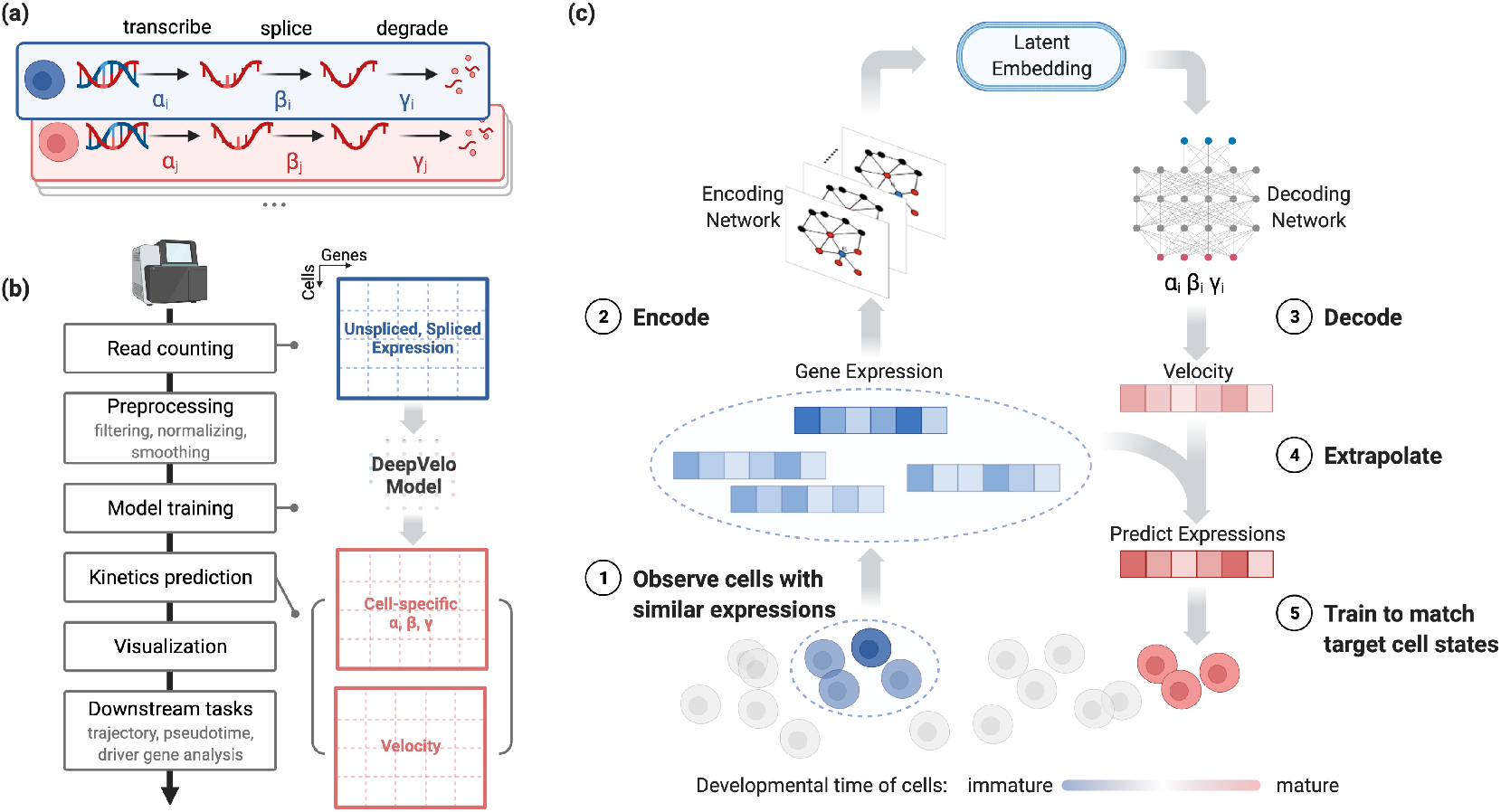
Overview of the DeepVelo pipeline and velocity prediction method. (a) DeepVelo estimates cell-specific transcription (*α_i_*), RNA splicing (*β_i_*) and RNA degradation rates (*γ_i_*). (b) Overview of the velocity analysis pipeline using DeepVelo. After read counting of unspliced and spliced mRNA, preprocessing is done to ensure the stability of model training (Online methods), followed by training and prediction of cell-specific kinetic parameters. These are used to estimate the RNA velocity and perform downstream analyses, such as visualization of velocity fields and pseudotime inference. (c) Overview of the DeepVelo neural network model. Query cells (dark blue) and similar cells (light blue) within a k-nearest neighborhood are input into the model. The Graph Convolutional Network (GCN) encoder module encodes their spliced/unspliced gene expression into latent space representations. The decoder module then predicts the kinetic rates for RNA velocity and extrapolates gene expression to future cell states. The model is optimized to match the extrapolation to observed cell states at later developmental stages. After training and optimization, these rates can be used to determine the RNA velocity vector for each cell through cell-specific rates of transcription, splicing and degradation.

DeepVelo models the kinetic rates per cell and per gene (Fig. 1a), providing sufficient expressive power for more faithful velocity estimates for individual cells. Given the unspliced gene counts *u*(*t*) and spliced gene counts *s*(*t*) for individual cells, DeepVelo estimates the derivatives of *s*(*t*) by modeling cell and gene-specific coefficients *α_i,g_*, *β_i,g_*, *γ_i,g_* using a deep neural network model (Fig. 1b,c). Specifically, we predict a cell’s velocity vector and extrapolate the cell state to match the future states extracted from the sequencing data (Fig. 1c). For each cell *i* in the population, we extract a group of neighbor cells 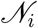 that have similar expression profiles. We take the profiles of cell *i* and neighbor set 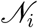 as the input to DeepVelo model. The model consists of stacked layers of GCNs and outputs the coefficients *α_i,g_, β_i,g_*, and *γ_i,g_* in the final layer. Using these coefficients, DeepVelo computes the velocity 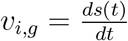 for each cell accordingly as in Eq.1.

To train the DeepVelo model, i.e. to update the parameters for accurate velocity prediction, we first extrapolate the cell state by adding the velocity derivative 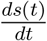 onto the original profile *s* (*t*). Then, DeepVelo computes the difference between the extrapolated state *s*(*t* + 1) and the real profiles of a group of downstream cells (The red cells in Fig. 1c). The DeepVelo model parameters are optimized to minimize this difference between the predicted future state and the actually observed ones (Online methods – 5.3). After sufficient training iterations, the model is finalized to provide accurate velocity estimates that take into account the transcriptional dynamics unique to individual cells.

We tested DeepVelo on a number of developmental datasets to determine RNA velocity, estimate cell-specific RNA kinetics, infer developmental pseudotime, and prioritize genes for their potential role in differentiation through driver gene estimation.

### 3.2 Recovering complex transcriptional dynamics for individual cells using DeepVelo

To test the ability to identify complex kinetics, we applied DeepVelo on a neurogenesis scRNA-seq dataset of the developing mouse dentate gyrus [11]. The data consists of tissue samples from two experimental timepoints, P12 and P35 (postnatal day 12 and 35), collected by a droplet-based single-cell RNA sequencing protocol (10x Genomics Chromium Single-Cell Kit V1). After pre-processing (Online Methods – 5.1), we calculated the RNA velocities using the proposed DeepVelo model and the dynamical model from scVelo [4]. The velocity plots are made by projecting the velocity vectors onto the UMAP [21]-based embedding of the data. In the velocity estimates (Fig. 2a), the granule cell lineage dominates the main structure, where the neuroblast cells develop into immature and mature granule cells. The directions of these velocity estimates between celltypes reflect the actual development orders [11].

**Figure 2:**
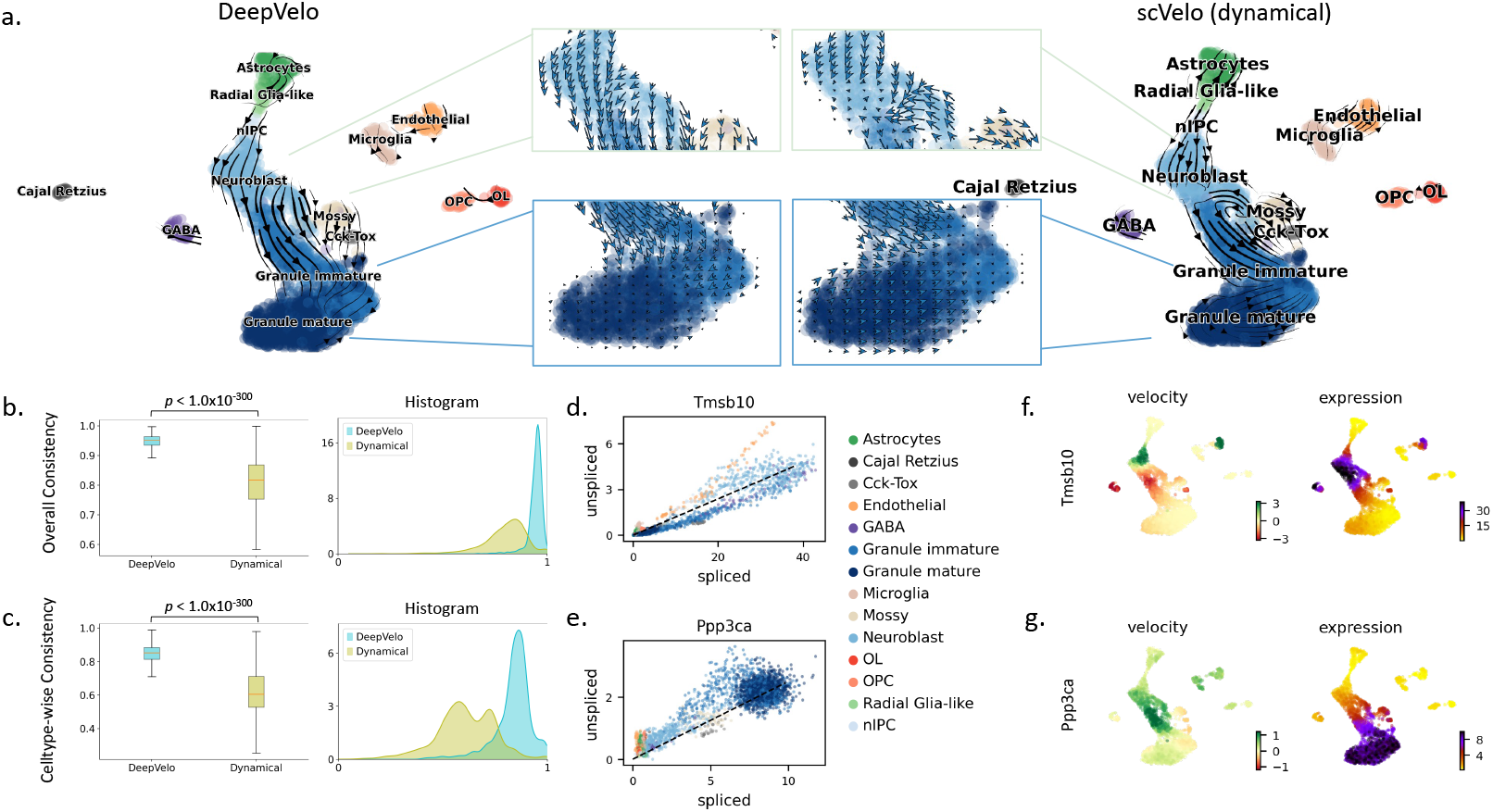
Fine-grained temporal patterns in neurogenesis predicted by DeepVelo. (a) Comparison of DeepVelo with the dynamical model from scVelo [4]. The direction and magnitude of velocities are projected as arrows onto the Uniform Manifold Approximation and Projection (UMAP) plot of gene expression values across cells. DeepVelo provides more consistent velocity estimates with respect to the developmental process from immature granule cells to mature granule cells. (b) The boxplot and histogram of the overall consistency scores for scVelo and DeepVelo, which indicate the consistency of velocity estimates in a local neighborhood of the data. (c) The box plot and histogram of the cluster/celltype-specific consistency scores, which utilize the neighborhood consistency metric on a per cluster/celltype basis. (d)(e) The spliced/unspliced phase portrait for *Tmsb10* and *Ppp3ca,* respectively. Celltypes are shown in the same color as in panel (b). (f)(g) Velocity and gene expression values projected onto UMAP plots for *Tmsb10* and *Ppp3ca,* respectively. Velocity and gene expression values show consistent patterns across celltypes: high velocity values (green in velocity plot) are correctly shown in the subset of cells developing to high gene expression values (purple in expression plot)

When examining the main lineage toward the terminal celltype of granule cells, although all models capture the principle direction, DeepVelo can show a more consistent flow from the neurogenic intermediate progenitor cells (nIPC) to neuroblasts, and finally to granule cells. DeepVelo particularly indicates that immature granule cells differentiate into mature granule cells in a manner more faithful to the true trajectory compared with the dynamical model (Fig. 2a – zoom-in panel).

The estimated velocities by DeepVelo show higher consistency in quantitative analysis. The consistency score is computed as follows – we first compute the average cosine similarity of the velocity vector of each cell to its neighbors, which is defined as the overall consistency. A similar neighbor-wise consistency was also proposed in scVelo [4]. However, the overall consistency could be biased toward over-smoothed estimations, which do not account for branching lineages. Therefore, we propose the cluster/celltype-wise consistency as a complement to the overall score, which computes the average cosine similarity of each cell’s velocity to all velocity vectors of the same celltype (Online Methods – 5.4). For both metrics, DeepVelo shows significant improvements over the scVelo dynamical method with significantly higher average consistency scores (Mann-Whitney U Test *p* < 1.0 × 10^-300^, Fig. 2b,c).

Examined at the individual gene level, DeepVelo shows biologically meaningful velocity patterns. For example, *Tmsb10* is one of the major regulators to the inferred dynamics of granule lineage, and it plays an important role in the development of hippocampal CA1 region [1]. In Fig.2f, velocities derived from the DeepVelo are consistent across velocities of neighboring cells. The region of cells showing high velocities of *Tmsb10* aligns well with the region of high *Tmsb10* expression. The same alignment is also observed in the example of another regulatory gene, *Ppp3ca* (Fig.2g). In further analysis (Fig.3a), we also observed that DeepVelo clearly disentangles the velocity vectors between the granule (blue) and endothelial lineages (orange), whereas, in the steady-state and dynamical models, both lineages have intertwined velocities. We discuss this advantage of celltype-specific prediction in Section.3.3.

**Figure 3:**
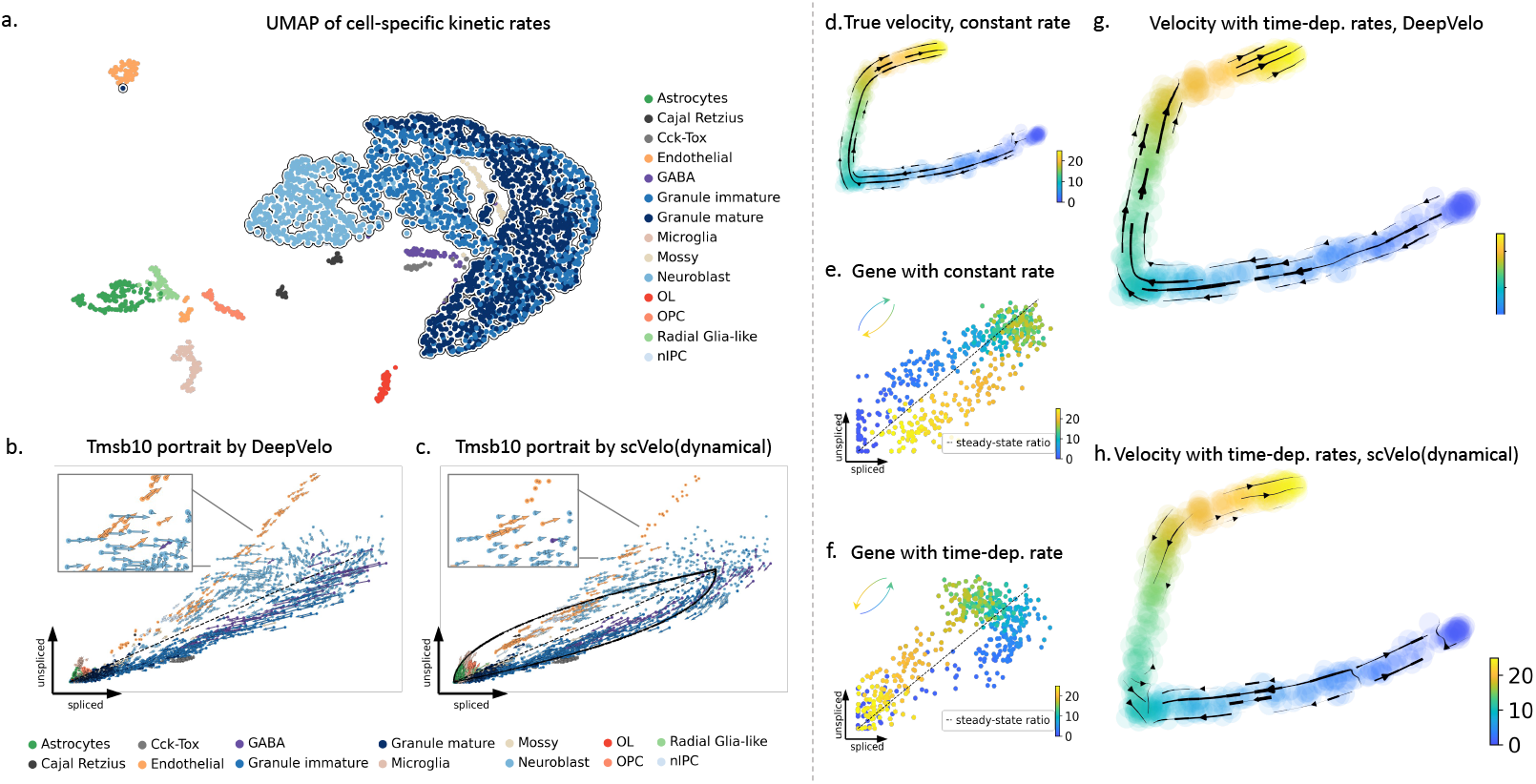
Velocity estimation for branching and time-dependent kinetic rates. (a) The UMAP projection of the estimated kinetic rates of 2930 cells in the dentate gyrus developmental data. Cells of the same celltypes are clustered together by kinetic rates. Further, cells from the same lineage (e.g. the outlined Granule lineage) are positioned closely. In general, the similarity of learned kinetic rates reflects the biological similarity of cells, although the DeepVelo model is unaware of celltype labels. (b) Projection of estimated velocity (arrows) onto the spliced/unspliced phase portrait of *Tmsb10* by DeepVelo. The endothelial cells undergo a separate trajectory on the phase portrait, aside from the main trajectory containing neuroblast cells, granule immature and granule mature cells. DeepVelo successfully captures both trajectories. In the zoomed view, cells within the same region comprising of different celltypes are correctly predicted to have distinct velocity directions. (c) Phase portrait of *Tmsb10* with RNA velocity predicted by the scVelo dynamical model. Only the main trajectory of granule lineage is captured, but the endothelial cells are predicted with incorrect directions. (d-h) A simulation of time-dependent degradation rates. The cell color indicates its pseudotime in simulation. (d) Reference velocity with constant kinetic rates. (e)(f) Constant and time-dependent degradation rates as shown on phase portraits. The gene with the time-dependent rate (f) undergoes a reversed trajectory. (g)(h) Estimated velocities by DeepVelo and scVelo, respectively, for the simulated 500 cells with time-dependent degradation rates. DeepVelo correctly recovers the directions from regions of earlier time to later ones.

### 3.3 DeepVelo’s cell-specific kinetic rates enable accurate quantification of time-dependent and multifaceted gene dynamics

Due to the cell-specific estimation of (*α_i,g_,β_i,g_,γ_i,g_* in Eq.1), DeepVelo for the first time provides a profile of individual kinetic rates for each cell. This enables new approaches for cell-specific trajectory analysis, visualization, and characterization. We show the UMAP projection of all cell-specific kinetic rates of 2930 cells (Fig.3a). Although DeepVelo is **unaware of the celltypes during training**, the learned kinetic rates are naturally clustered into groups aligned with celltypes. Further, clusters of cells from the same lineage (e.g. the outlined granule lineage) are positioned closely compared to other cells. Overall, the similarity of learned kinetic rates reflects the biological similarity of cells at both the celltype and lineage levels. This indicates that DeepVelo can estimate kinetics that reflects the dynamics of individual cells as opposed to the entire dataset.

Velocity-associated kinetic rates across cells may vary for genes undergoing dynamic regulation involving multiple processes. For example Battich, Stoeger, and Pelkmans [3] observed varying kinetic rates in the differentiation of intestinal stem cells. These varying kinetics are often misinterpreted in existing velocity methods [5]. This stems from the fact that the kinetic rates in previous methods are modeled as constant cell-agnostic coefficients in first-order equations (Eq.2), which lack the ability to model multifaceted dynamical variation. In contrast, DeepVelo estimates transcriptional dynamics for different celltypes and cell states by introducing cell-specific kinetic rates, leading to better velocity estimation in time-dependent and complex multi-lineage systems. Here, we show this improvement using two challenging scenarios:

#### (1) Estimating velocity for genes that are separately regulated in two lineages

We used the previously analyzed dentate gyrus cell population and determined genes with multifaceted kinetics [11]. *Tmsb10* shows multiple kinetic regimes and undergoes multiple trajectories. We plot the spliced and unspliced reads across all cells in this dataset, in other words, the phase portrait of *Tmsb10* (Fig.2d). The cells in the granule lineage (including neuroblast, granule immature and granule mature celltypes) form a cyclic trajectory. Meanwhile, the endothelial cells are not a part of the granule lineage and undergo a separate trajectory. These two regimes are likely regulated by different kinetic rates.

DeepVelo correctly predicted the RNA velocity patterns for both regimes (Fig.3b). For the granule lineage, DeepVelo captures the direction of velocity from neuroblast cells to granule immature cells and then to granule mature cells. For the endothelial cells, the predicted velocity direction correctly points to the position of the same celltype with amplified spliced reads. We also found that DeepVelo learns to assign similar velocity directions for cells of the same type. In contrast to DeepVelo, scVelo forces the RNA velocity to follow the cyclic trajectory assumed by the model (Fig.3c). As a result, although scVelo successfully captures the trajectory for the granule lineage, it incorrectly points the velocity estimates of endothelial cells to the position of neuroblasts (Fig.3c – Zoom-in panel).

Additionally, DeepVelo is capable of predicting distinct velocity directions for cells within the same region (Fig.3b). The cells in the zoomed view, including both the endothelial and neuroblast cells, employ similar RNA dynamics (through the levels of spliced and unspliced reads) of *Tmsb10.* However, the distinct directions for each celltype are correctly predicted by DeepVelo. This is due to the ability of DeepVelo to estimate distinct sets of kinetic rates across celltypes, as shown in Fig.3a. In contrast, scVelo uses constant kinetic rates per gene and predicts a uniform direction for the same region of cells. Overall, a cell-specific model such as DeepVelo broadens the application of RNA velocity for genes with multifaceted kinetics, such as *Tmsb10* in the dentate gyrus developmental data.

#### (2) Estimating velocity for genes with time-dependent kinetic rates

We simulated a population of 500 cells and 30 genes using the simulator provided by the scVelo package [4]. We first determined the reference velocity in the setting of constant kinetic rates across cells (Fig.3d,e). From here, the degradation rate, *gamma,* of 3 out of 30 genes was set to increase over time. As a result, the genes underwent a reversed trajectory as shown in the respective phase portrait (Fig.3f). This simulation procedure of reversed dynamics was originally proposed in Bergen et al. [5], and it sets up a challenging scenario for the estimation of RNA velocity. The resulting velocity plots of DeepVelo and the dynamical model of scVelo are shown in Fig.3g,h, and scVelo struggles to predict velocities from early to later timepoints while DeepVelo is able to recover the correct velocity directions from regions of earlier to later pseudotime. This advantage is because DeepVelo learns to find potential future cell states by integrating across all genes (Online Methods – 5.3), thus it is more robust to the time-reversed directions of a portion of genes in the dataset.

### 3.4 Tracking the ordering of cellular development using DeepVelo and diffusion pseudotime

The RNA velocity estimated by DeepVelo can also be used to improve the prediction of pseudotime for cell states across a developmental trajectory. We first compute the velocity connectivity graph to represent cell-cell relationships and use this graph as the basis to compute a diffusion pseudotime [10] mapping (Online Methods – 5.5). We compare the pseudotime estimates (Fig.4a) using DeepVelo with the latent time (Fig.4c) estimates by the dynamical model from scVelo on a scRNA-seq dataset of pancreatic endocrinogenesis with ground-truth temporal measurements. For the velocity plots, DeepVelo successfully demonstrates the main structure of EP cells developing into terminal celltypes – alpha, beta and delta (Fig.4b) with more consistent velocities (Fig.4e). For pseudotime comparison, both methods provide accurate predictions. Notably, DeepVelo more faithfully preserves the time ordering between the terminal states of Alpha and Beta cells (Fig.4a), where the Alpha cells are developed earlier at E12.5 and the Beta cells appear later at E15.5 (Fig.4f,g).

**Figure 4:**
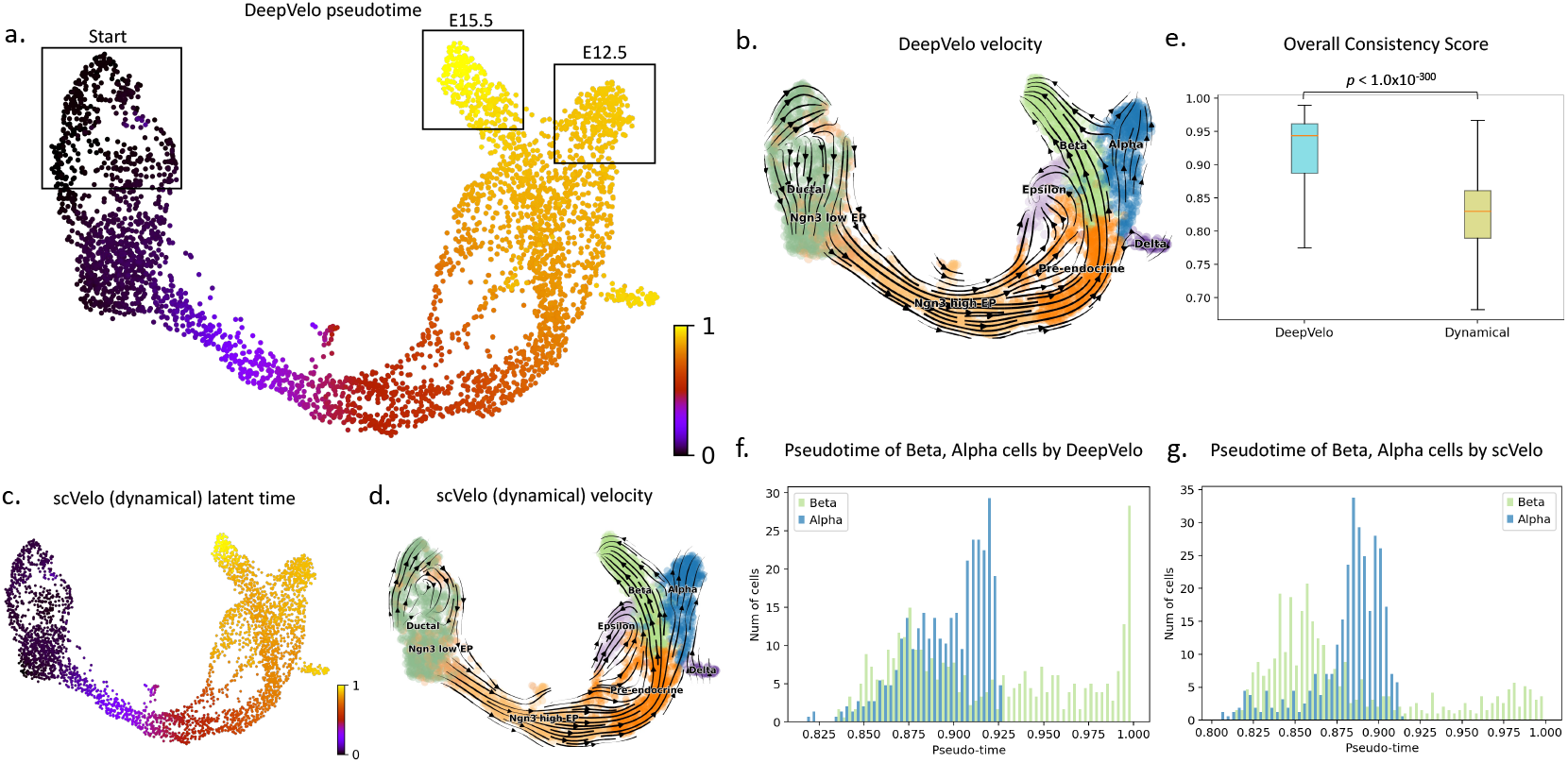
Velocity and pseudotime plots for pancreatic endocrinogenesis [2]. (a) The pseudotime prediction from DeepVelo accurately assigns alpha and beta cells to accurate developmental timepoints. Particularly, the progenitor cell cluster is correctly located at the upper left quadrant of the UMAP projection. The difference between the terminal alpha and beta cells is well captured, where alpha cells were developed earlier at E12.5 and beta cells appeared later until E15.5. (b) Velocity values derived from DeepVelo are projected onto the UMAP-based embedding and visualized. DeepVelo successfully captures the main structure of EP cells developing into the terminal celltypes of alpha, beta and delta cells. (c),(d) For comparison, the latent time and velocity computed by the dynamical model from scVelo. (e) Distribution of the overall RNA velocity consistency scores for DeepVelo and scVelo. (f,g) The histogram of pseudotime predictions for beta and alpha cells, by DeepVelo and the scVelo dynamical model, respectively. Beta cells are expected to have a larger percentage of cells with higher pseudotime values, which is true of the DeepVelo predicted values.

### 3.5 DeepVelo infers functionally relevant lineage-specific genes and processes in hindbrain development

To test velocity methods in a complex setting with multiple lineages, we applied methods to a mouse hindbrain development dataset [30] (Fig.5a). Specifically, we filtered the data corresponding to the junction and differentiation between the GABAergic and gliogenic lineages (Online Methods – 5.1). In a multi-faceted system such as this, which is typical of developmental scRNA-seq datasets, considering cell-agnostic kinetic rates is haphazard because of the different RNA velocity dynamics among lineages. DeepVelo’s ability to learn cell-specific kinetic rates alleviates this assumption and accounts for the multi-faceted differentiation of the GABAergic and gliogenic lineages and their respective celltypes. The result of DeepVelo (Fig.5b) shows the RNA velocity over the developmental process from Neural stem cells to the differentiating GABA interneurons and gliogenic progenitors. We performed trajectory inference using directional PAGA [32] over the velocity graph of DeepVelo. We found that DeepVelo was able to recapitulate ground-truth differentiation patterns – specifically the branching between VZ progenitors and differentiation GABA interneurons and gliogenic progenitors (Fig.5c). The cluster of neural stem cells is well recognized as the origin celltype with outward velocity arrows and a low pseudotime index, while the GABA interneurons are confirmed as a terminal celltype with incoming velocity arrows and a high pseudotime index. In comparison, the scVelo dynamical model predicts partially inverse velocity directions for the gliogenic progenitors, leading to incorrect relations in the inferred trajectory (Fig.5f, highlighted regions).

**Figure 5.**
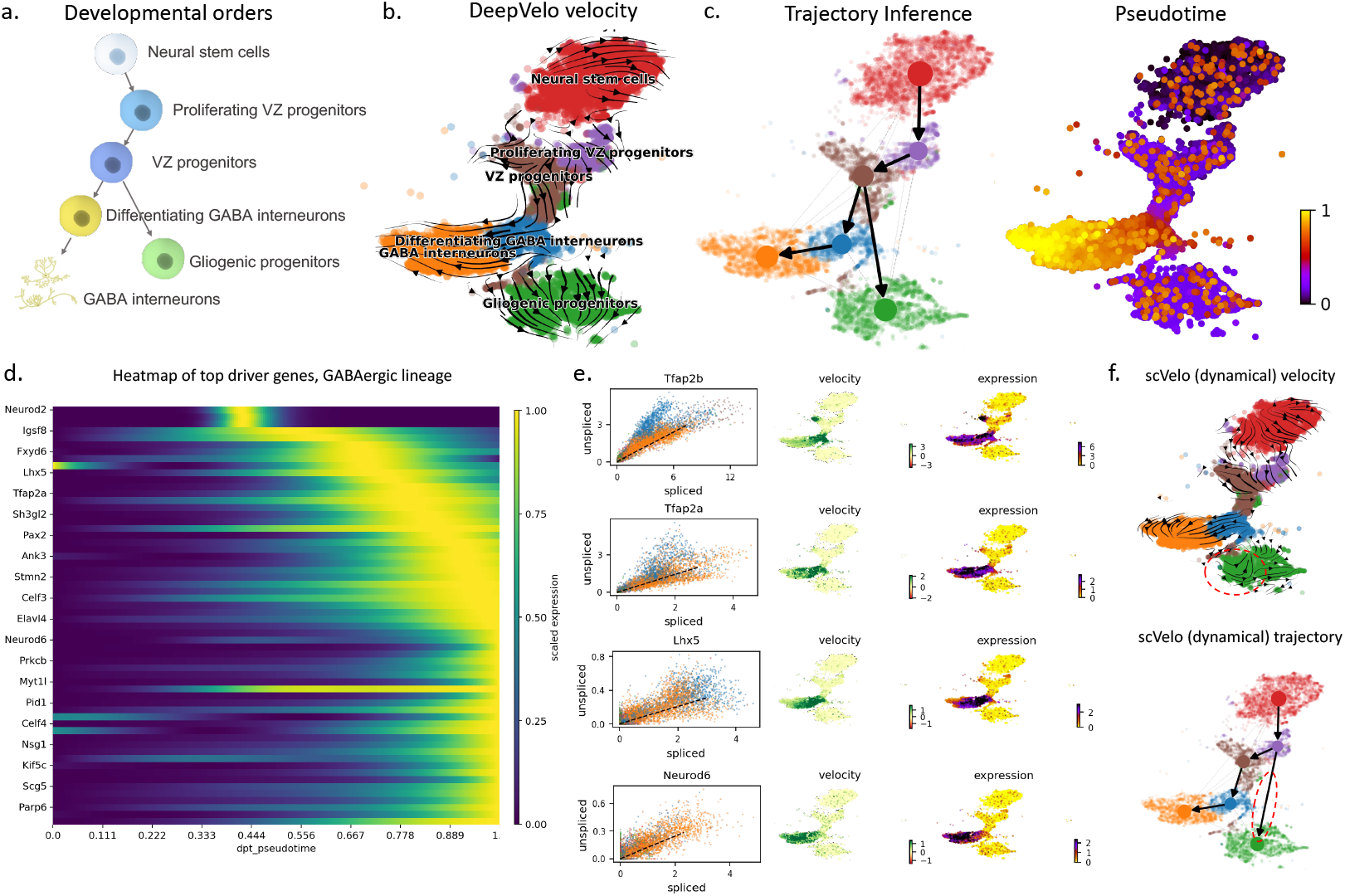
Velocity, trajectory, and driver gene estimation of developing mouse hindbrain cells. (a) The putative developmental order for six celltypes in early mouse hindbrain development. (b) The velocity projected onto the t-distributed stochastic neighbor embedding (tsne) plot of gene expression. DeepVelo’s RNA velocity reveals the temporal order in the developing mouse hindbrain, including cells from early progenitors, GABAergic, and gliogenic lineages. (c) Velocity-based PAGA trajectory inference using DeepVelo’s velocity estimates. The predicted trajectory correctly reflects the developmental relations shown in (a). (d) The top 60 driver genes with highest correlation to the GABAergic lineage computed using DeepVelo’s velocity estimation. The horizontal coordinates represent the pseudotime estimates. (e) Gene phase portrait, velocity, and gene expression plots of selected driver genes. Known functional genes in the GABAergic lineage – *Tfap2b, Tfap2a, Lhx5*, and *Neurod6* – are computed among the top driver genes. (f) The velocity plot and trjectory inference using the scVelo dynamical model.

Using the velocity vector for each cell, we built a connectivity graph (Online Methods – 5.5) of the scRNA-seq data. CellRank [18] is a recent visualization and analysis toolbox for scRNA-seq data that utilizes the connectivity graph to predict cell’s fate mapping, which corresponds to the probability of the cell differentiating to a terminal state in the lineage(s). After determining cell fate, gene importance for differentiation can be calculated based on the correlation of gene expression with transition and differentiation probabilities towards all terminal states. The genes that display dynamical behavior across a lineage are termed putative ‘‘driver genes”, as these are the genes most likely to be involved in regulating the differentiation process itself. CellRank has been reported to work well with other velocity methods, such as scVelo, to infer lineage-specific drivers. We incorporated this toolbox with the predicted velocity connectivity graph from DeepVelo and determined driver genes in the variable gene subset of the data for both the GABAergic and gliogenic lineages.

Within the top 100 driver genes across both lineages of interest, we observed groups of genes showing particular abundance in specific celltypes in a temporal manner (Fig.5d). For example, *Tfap2a, Tfap2b,* and *Lhx5,* which are two known differentiation genes involved in the specification of GABAergic interneurons during hindbrain development, are listed in the top 100 driver genes from DeepVelo for the GABAergic lineage (Fig.5e) [34, 23]. Similar results were found for the gliogenic lineage from DeepVelo, with detection of known glial cell differentiation regulators in *Hes1* and *Sox9* (Supplementary Table 1) [33, 31]. DeepVelo also picked up hits that were novel and not detected by scVelo, such as *Neurod6* in the GABAergic developmental lineage (Fig.5e). Although the role of *Neurod6* in the differentiating GABAergic interneurons and their development is unclear, previous literature has indicated the gene’s involvement in regulating the specification of inhibitory GABAergic interneuron subpopulations in the hindbrain and spinal cord [27]. This indicates a testable link and hypothesis for the differentiation of these cells in the junction within the GABAergic and gliogenic lineages, highlighting the ability of DeepVelo to guide searches of functional genes in scRNA-seq data and potential drivers of the differentiation process.

To compare the results of driver analysis when employing CellRank with different velocity outputs, we determined driver genes for the gliogenic and GABAergic lineages using both scVelo and DeepVelo. As the complete set of genes driving differentiation in the complex hindbrain developmental system is unknown, we sought to infer the relevance of inferred driver genes in two ways: 1) By considering their overlap with predicted marker genes from the original analysis, as these genes are characteristic of celltype identity and should be correlated with lineage specification, and 2) By considering their overlap with transcription factors (TFs), as TFs are the main elements responsible for differentiation and establishing transcriptional and cellular identity. We analyzed and compared the top 100 driver genes for both the GABAergic and gliogenic lineages predicted by the scVelo and DeepVelo methods (Supplementary Table 1). DeepVelo predicted driver genes that overlapped with more of the original markers from Vladoiu et al. [30], for both the GABAergic and gliogenic lineages (Fig.6a) (Supplementary Table 2). Further, to determine the signal across all driver genes, not limiting to the top 100, we determined the rankings of known marker genes in the GABAergic and gliogenic lineages across all tested driver genes. These rankings were determined based on the correlation scores, which indicate the relative importance of driver genes to a specific lineage. In this case, DeepVelo had higher rankings compared to scVelo for known GABAergic marker genes in the driver analysis (Mann-Whitney U Test *p* = 1.376 × 10^-07^), while the ranking differences in the gliogenic lineage were non-significant (Fig.6b). When examining the transcription factor overlap in the top 100 driver genes, DeepVelo had more hits than scVelo for the GABAergic lineage, and an equal number of hits for the gliogenic lineage (Fig.6c).

**Figure 6:**
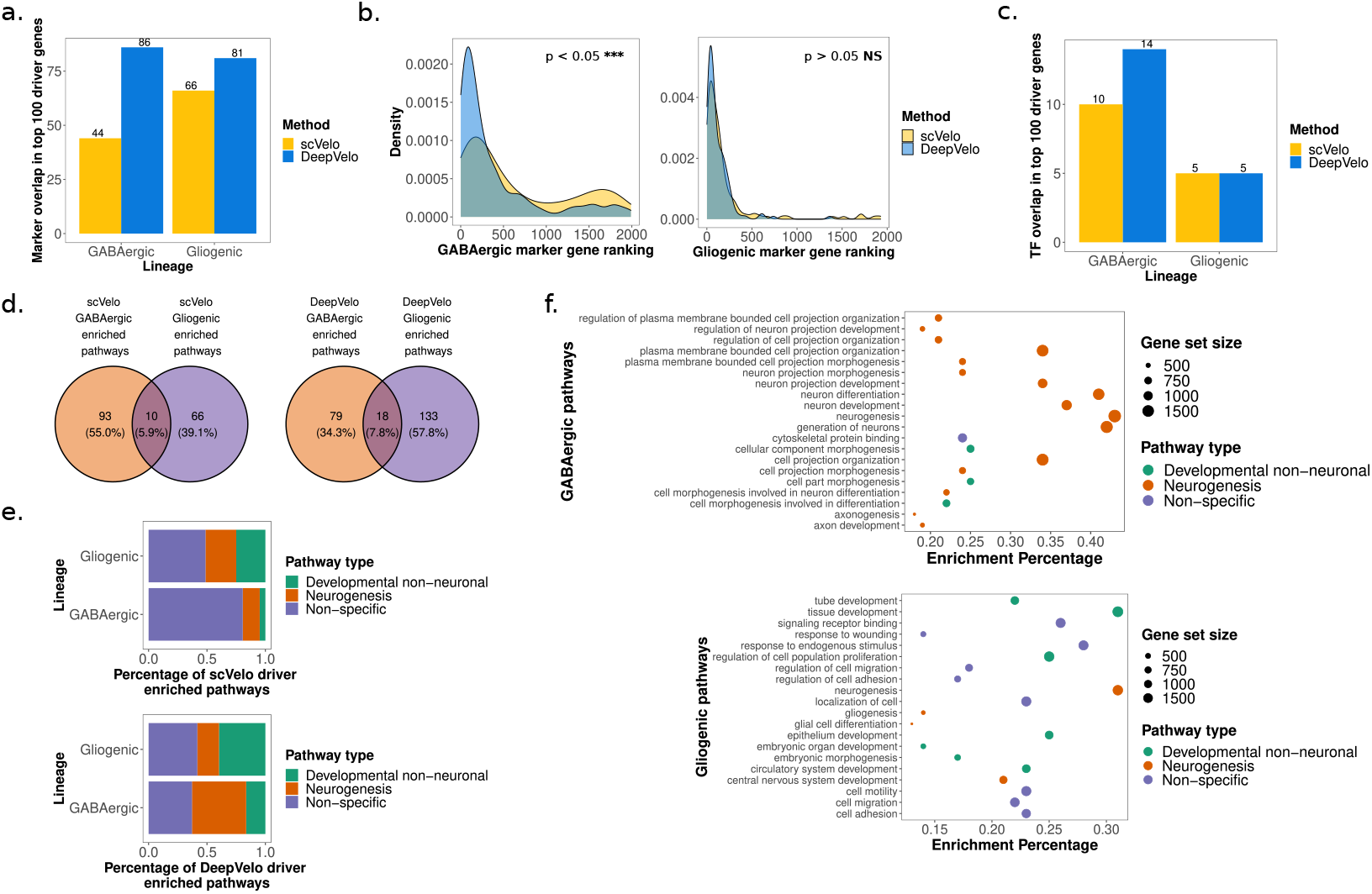
Functional enrichment of DeepVelo predicted driver genes. (a) Overlap of the top 100 driver genes from scVelo and DeepVelo for GABAergic and gliogenic lineages with annotated lineage marker genes. (b) Ranking density of marker-overlapping driver genes (across all 2000 tested genes) for scVelo and DeepVelo, separated by the GABAergic and gliogenic lineages, respectively. (c) Overlap of top 100 driver genes from DeepVelo and scVelo for both lineages with annotated transcription factors. (d) Pathway enrichment analysis results for the top 100 scVelo and DeepVelo driver genes, respectively, in the GABAergic and gliogenic lineages. (e) Functional signal in the enriched pathways for scVelo and DeepVelo, based on the presence of pathways involved directly in neurogenesis (“Neurogenesis”), not specific to neurogenesis but involved in development (“Developmental non-neuronal”), and not specific to either development or neurogenesis (“Non-specific”). (f) The top 20 DeepVelo pathway enrichment analysis results, based on FDR corrected *p*-values, for the GABAergic and gliogenic lineages, respectively.

For further examination of the results of driver analysis, we took the top 100 driver genes for the GABAergic and gliogenic lineages from DeepVelo and sought to determine their functional signal as gene-sets through pathway enrichment analysis. Overall, 97 and 151 pathways were found to be significantly enriched for the GABAergic and gliogenic lineages, respectively, for DeepVelo (Fig.6d) (Supplementary Table 3). These pathways were analyzed for the presence of neurogenesis and developmental results, for which we did see a functional enrichment in both lineages (Fig.6e). More specifically, the top 20 pathways for each lineage, ranked in terms of FDR-corrected p values, revealed enrichment of pathways relevant to neuronal differentiation processes (Fig.6f). In the GABAergic lineage, enriched pathways included: *regulation of neuron projection development, neuron differentiation,* and *neurogenesis* (Fig.6f). The results from the gliogenic lineage had an even more relevant terms, namely *gliogenesis* and *glial cell differentiation* (Fig.6f). When comparing these results with pathway analysis performed on the scVelo top 100 driver genes, we observed a much lower percentage of functional enrichment for neurogenesis and developmental pathways compared to DeepVelo for the GABAergic lineage (Fisher’s Exact Test p = 1.407 × 10^-09^) (Fig.6e), while the difference between the gliogenic results was non-significant. These functional pathway enrichment results highlight the relevance of the driver genes predicted by the DeepVelo method and increased functional relevance compared to those predicted by scVelo.

### 3.6 DeepVelo is computationally robust and efficient across multiple scRNA-seq datasets

To examine the robustness of the DeepVelo RNA velocity estimates across settings, we tested DeepVelo on five different scRNA-seq datasets. Apart from the previously analyzed datasets, DeepVelo also recovers accurate RNA velocity vectors and developmental relations on a large-scale dentate gyrus data from La Manno et al. [16] (Supplementary Fig.S3). On all tested datasets, DeepVelo achieves higher average scores and lower variance in terms of the overall consistency compared to the scVelo dynamical model and the scVelo stochastic model (Supplementary Table 4).

We further visualized the influence of key hyperparameters – including the GCN layer size, gradient descent learning rate, and number of training epochs – on the dentate gyrus neurogenesis data (Supplementary Fig.S4). DeepVelo is robust to changes in these hyperparameters and consistently estimates the biologically accurate RNA velocity directions, especially for the main granule lineage.

Lastly, we compared the computational runtime of DeepVelo with other velocity estimation methods. Using the same CPU (central processing unit) device, DeepVelo (cpu) achieved a 4 fold speedup with respect to the scVelo dynamical model. Using a more powerful GPU (graphical processing unit) for the deep learning backbone, DeepVelo (cpu+gpu) can be further accelerated 10-20 times across datasets. For example, DeepVelo completed the training and estimation for the 13501 cells of developmental hindbrain data in just 36 seconds (Supplementary Fig.S5).

## 4 Discussion

DeepVelo offers a novel velocity estimation framework that goes beyond assumptions of constant RNA splicing and degradation rates, and instead estimates these rates at a cell-specific level. By analyzing the performance of DeepVelo and existing velocity estimation techniques, we have demonstrated that DeepVelo’s cell-specific estimation through a novel deep learning method allows for the detection and specification of multiple lineages in calculating RNA velocity. Realistic single-cell RNA sequencing settings will likely have more than one lineage/trajectory in a given sample, and thus it is imperative to develop methods that can account for these multifaceted dynamical systems. DeepVelo’s ability to model these multifaceted dynamics was demonstrated through analysis of complex differentiation systems, such as the development of the dentate gyrus, pancreatic endocrinogenesis, and the hindbrain development. Lastly, we demonstrated that DeepVelo can be utilized to identify functionally relevant genes that are enriched along the differentiation trajectory. We envision that DeepVelo will be more readily applicable to these realistic developmental settings as compared to previous techniques.

DeepVelo internally predicts the first-order derivative of expression per gene based on the transcriptome-wide reads of all selected genes. The ability to learn the interaction/regulation between genes could be further explored, for example, by replacing the GCN model with recent transformer networks [29] which could explicitly model the interaction of internal gene representations. This could allow for more interpretable velocity and driver-gene estimates, by considering correlations of kinetics and expression patterns between genes and cells. Recent work shows promising research directions by extending the velocity of cellular dynamics from RNA to proteins [8], epigenomics [26], and multi-omics velocities [20]. DeepVelo could be naturally updated and well fitted into these settings by enriching input and output space with additional-omics information. Ultimately, the estimation of cell-specific kinetics across multiple steps in the central dogma may increase the signal-to-noise ratio [5] and further accurately capture information related to cellular development.

A major limitation of driver analysis through RNA velocity estimation is potentially spurious driver genes being picked up due to the correlation of gene expression during differentiation. Although key regulators will display dynamical expression behavior during lineage specification, the same is likely to be true of their downstream targets and other “passenger” genes, resulting in high likelihoods towards being a driver/regulator. This is likely the reason why a significant transcription factor enrichment was not observed in the top 100 driver genes in the hindbrain developmental data for either scVelo or DeepVelo. We envision a more comprehensive driver analysis technique would take into account multi-lineage probabilities (preventing negative correlation between top drivers of two lineages) and would factor in correlations between driver analysis results, thereby removing spurious hits. Apart from the driver gene analysis, building up a theorem to verify the confidence of velocity estimation is another challenge. Empirical metrics, such as the consistency of velocity directions among neighbor cells, have been used in existing techniques [4, 24]. However, there is a lack of probabilistic tools to test the kinetics estimated by either previous methods or DeepVelo. We leave this to future works.

RNA velocity techniques have allowed for insights into biological differentiation from single-cell RNA sequencing data that go beyond oversimplified trajectory inference models, and instead infer dynamic processes that indicate the direction and magnitude of differentiation potential. Although many major limitations and assumptions for RNA velocity methods still exist, we anticipate that continued methodological development in this field will lead to better tools to study differentiation and development in a single-cell setting. DeepVelo overcomes limitations of previous techniques in a major aspect with regards to cell-specific model estimates, and can be used for more robust velocity estimation in multi-lineage systems, yielding better biological insights into real and complex developmental systems.

## 5 Online methods

### 5.1 Preprocessing the scRNA-seq data for DeepVelo

The dentate gyrus neurogenesis [11] and pancreatic endocrinogenesis [2] data are available at the National Center for Biotechnology Information’s Gene Expression Omnibus repository. The accession number is GSE95753 and GSE132188. In this work, we use the zipped data of these two sequencing datasets provided by the scVelo packageBergen et al. [4](https://scVelo.org). The data is in h5py file format and contains spliced and unspliced gene readout.

Mouse hindbrain developmental data from Vladoiu et al. [30] was used to test velocity techniques for estimation at a lineage junction. As the data was not available in loom format for velocity analysis, fastq files were reprocessed into loom files using kallisto reference-free alignment through the loompy pipeline [6]. This was done individually for each timepoint (E10, E12, E14, E16, E18, P0, P5, P7, P14) and processed loom files were concatenated. For the purposes of the analysis, the junction between the GABAergic and gliogenic lineages was utilized. The following celltypes were subset from timepoints E10, E12, E14, E16, E18, P0, P5, P7, and P14 – Neural stem cells, Proliferating VZ progenitors, VZ progenitors, Differentiating GABA interneurons, gliogenic progenitors, and GABA interneurons. Estimates of spliced and unspliced counts from the kallisto quantification method were used for testing DeepVelo and scVelo.

Processing of unspliced and spliced counts in differing formats was done via three steps and using the scVelo package. First, the spliced and unspliced gene matrices were normalized across genes. In more detail, preprocessing includes expression matrix normalization and nearest-neighbor-based smoothing. We used the `scv.pp.filter_and_normalize` function from scVelo for these steps with default parameters. We selected the top 2000 genes with the most spliced and unspliced gene counts across cells. The principal components are computed afterward using logarithmized spliced counts, and then we smooth the expression reads using the average of 30-nearest-neighbors for each cell.

### 5.2 Modeling individual transcriptional dynamics

The transcriptional dynamics depicts the process from generation to degradation of mRNA molecules. It captures unspliced premature mRNAs *u*(*t*) with transcription rate *α*, splicing into mature mRNAs *s*(*t*) with rate *β* and the degradation of spliced mRNA with rate *γ*. The simplified gene-specific dynamics with constant splicing and degradation rates are

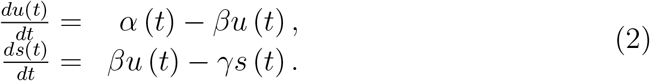

This equation is used in existing velocity estimation methods, and it omits the difference in kinetic rates (*α,β,γ*) across celltypes. Instead, we propose a new deep learning method to capture individual cell kinetics.

First, we build a graph convolutional network model to predict cell-specific kinetic rates. In this work, we employ a nearest neighbor graph based on the gene expression of all sequenced cells 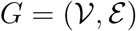. The vertex 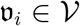 in the graph denotes the expression reads of a cell *i*, which include its spliced and unspliced gene expression 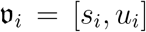. A cell *i* is connected to cell *j* (i.e. 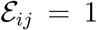) if cell *j* is the one of top 30 nearest neighbors based on the Euclidean distance of the gene expression. We input this neighbor graph to DeepVelo. We chose the graph representation because it considers the vicinity of local cells’ gene expression. This has more expressive power than the expression of individual cells because of the sparse and noisy nature of sequenced reads. Taking the neighborhood expression into account smooths the velocity estimation.

Graph convolutional network (GCN) is a type of deep neural networks that learns node embeddings based on message passing along the graph edges [15]. Given a graph with nodes 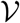 and adjacency matrix *A*, a multi-layer neural network is constructed on the graph with the following layer-wise propagation rule:

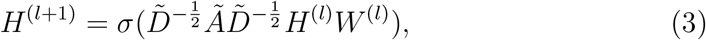

where *H*^(*l*)^ denotes the node feature vectors at the *l*-th layer, *Ã* = *A* + *I_N_* is the adjacency matrix with self-connections, 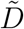 is the diagonal degree matrix such that 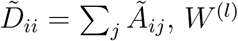 is the layer-specific trainable parameter matrix, and *σ* is the RELU activation function.

In this work, the input feature 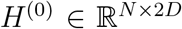 to GCN is the cellular gene matrix. Each row in H stands for the aforementioned vertex 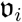. *H* contains the population of *N* cells, and the dimension 2*D* equals the number of selected spliced and unspliced genes combined, *D* = 2000 by default. The adjacency matrix *A* ∈ ℝ^*N×N*^ depicts the aforementioned nearest neighbor graph, where the element at position *i,j* has value 1 if the cell *j* is one of the nearest neighbors of cell *i*, otherwise the value is 0. The GCN model consists of stacked graph convolution layers, i.e. Eq. 3. The output of the final layer *H^L^* is processed by a fully connected neural network, which then yields the estimated velocity parameters *α* ∈ ℝ^*N×D*^, *β* ∈ ℝ^*N×D*^ and *γ* ∈ ℝ^*N×D*^ for all cells and genes.

Finally, the estimated velocity 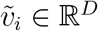 for each cell is computed as

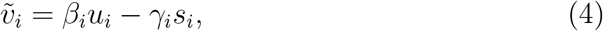

where *β_i_* and *γ_i_* are the *i*-th row in *β* and *γ*, *u_i_* and *s_i_* are the unspliced and spliced reads of cell *i*.

DeepVelo also supports estimation of the derivative of unspliced RNA, namely 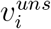, which is an estimation for the 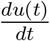 in Eq.1.

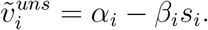

### 5.3 Probabilistic learning for RNA velocity

In this section, we propose a probabilistic learning framework for RNA velocity to optimize the velocity prediction in Eq.4, and then introduce the specific training objective following this framework.

#### 5.3.1 Extrapolate cell states from a probabilistic perspective

The RNA velocity is defined as the time derivative of spliced mRNA (Eq. 1). For a specific cell *i* out of the sequenced cell population Ω, the velocity vector *v_i_* contains the derivative for all genes, as

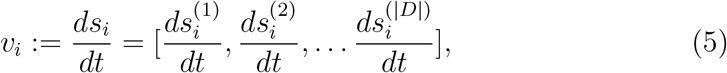

where 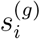 denotes the amount of spliced mRNA of one gene. *s_i_* is the spliced gene expression vector containing 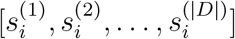.

We introduce the multivariate random variable *G*_*i,τ*(*i*)_ to represent the (spliced) gene expression that a cell *i* could have at its developmental time *τ*(*i*), where *τ* is an operator to obtain a cell’s current time in its developmental process. Thus, the scRNA-seq results could be viewed as an observation of *G*_*i,τ*(*i*)_ taking the value *s_i_*. For simplicity, let us use *t* = *τ*(*i*) as the time of cell *i*. Similarly, we define the random vector *V_i,t_* as the possible velocities that cell *i* can take at time *t*, and *v_i_* is an observation of *V_i,t_*. The relation between the expression and velocity random vectors is,

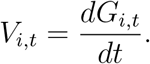

We can use the forward difference to approximate the derivative if the time interval is sufficiently small, as

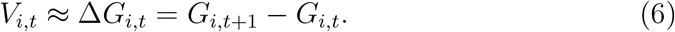

Notably, it is impossible to directly observe the future stage *G*_*i,t*+1_ for cell expression from scRNA-seq, because the sequencing protocol is destructive, and the cells no longer exist after sequencing. Thus, the estimation of *G*_*i;t*+1_ is required.

DeepVelo utilized the mRNA expression of developmentally close-by cells to estimate *G*_*i;t*+1_ and for this reason, we introduce the **continuity assumption**: we assume that the sequencing data captures a continuous spectrum of cells in consecutive development stages. Particularly, there exists a *t* + 1 neighborhood, 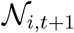, in the sequenced cell population, so that the gene expression of these cells within the neighborhood are similar enough to the potential expression of cell *i* at *t* + 1. In other words, the expected expression of the *t* + 1 neighbor cells have the same distribution as the expression of cell i at *t* +1. In comparison to the previous strict assumptions (i.e. the observation of steady states or the global constant kinetic rates) in existing approaches, the continuity is primarily satisfied in sequencing data of large cell populations. Formally, the continuity can be expressed as

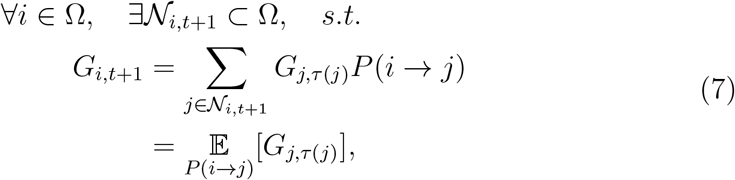

where *i* → *j* denotes that cell *i* develops at time *t* + 1 into a cell that has the same gene expression vector as cell *j*, and *P*(*i* → *j*) is the probability of this event. The expectation of *G*_*i,t*+1_ over all cells in the sequenced population Ω is

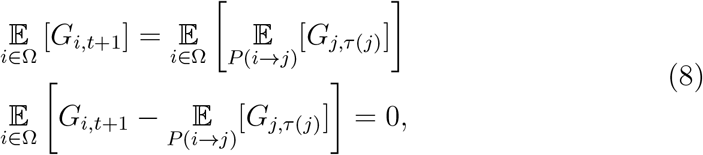

Taking in Eq.6, we have

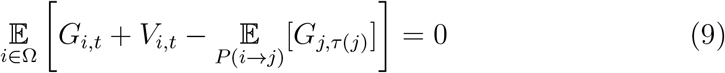

The observed sequenced expression in a large cell population can be used to derive the Monte Carlo estimation of the outer expectation over cell *i*. Assume each cell expression vector si is sequenced independently,

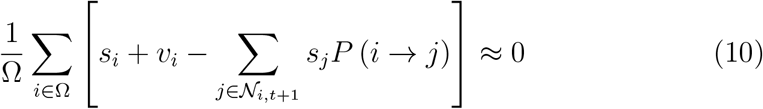

Because the *v_i_* and 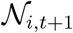 are not directly observed, given a set of estimated 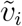 and 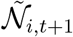, we use the (gene-wise) squared difference as an objective to measure how close to zero the value in Eq.10 is.

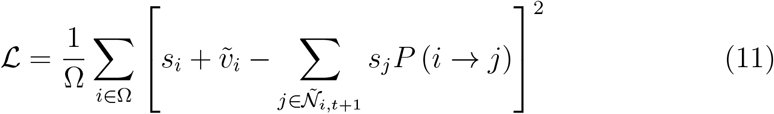

This equation provides a general objective for any RNA velocity methods that generate the estimation of 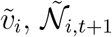 and *P*(*i* → *j*).

#### 5.3.2 Training the DeepVelo model

We follow the Eq.11 to develop the objective to optimize the parameters of DeepVelo model. The objective computes the difference between the estimated velocity 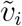 (Eq.4) of DeepVelo and possible future cell states.

We first select *K_c_* number of nearest neighbor cells for each cell *i* by computing the pairwise distances of spliced gene expression. By default, we compute the Euclidean distance of the first 30 PCA dimensions of gene expression vectors. These selected cells compose the neighborhood of cell *i*, i.e. 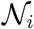. We estimate the *P*(*i* → *j*) using

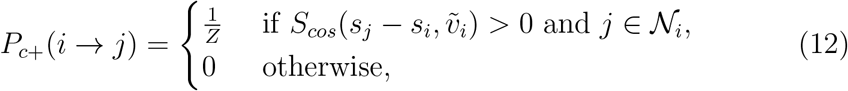

where *S_cos_* denotes the cosine similarity and *Z* is a normalizing factor, i.e. *Z* equals to number of cells in 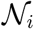 satisfying 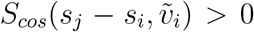. The intuition of *P*_*c*+_ is that if the sequenced data satisfy the continuity assumption and the time interval between *t* and *t* + 1 is small enough, then the possible future cell state 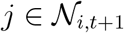 is also close to the cell state of current cell i. Therefore, given a sufficient large 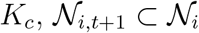. Further in Eq.12, We use the cosine similarity between the estimated velocity 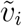 and the expression difference *s_j_* — *s_i_* to select the possible target cell *j* that aligns with the velocity direction.

Notably, the Eq.6 is the forward difference operation. Similarly, we can also include the backward difference *V_i,t_* = *G_i;t_* — *G*_*i,t*-1_ and project the cell *i* into *t* — 1. We first compute the probability of cell *i* developed from cell *j, P*_*c*-_ (*i* ← *j*), as follows

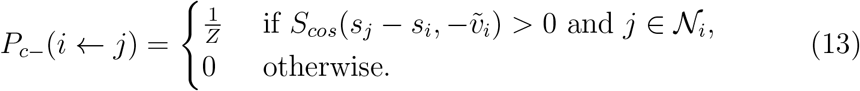

We then used this in the computation of 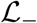 in Eq.14. The sum of 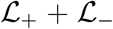 is symmetric to either 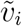 or 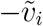, which creates a challenge to determine the correct velocity direction. To resolve this issue, we know from Eq.1 that the velocity across cells should be positively correlated to the unspliced expression, *u_i_*, and negatively correlated to the spliced, *s_i_*. We add the Pearson correlation in Eq.14 term 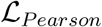 to promote the correct direction. The aforementioned objective terms are as follows

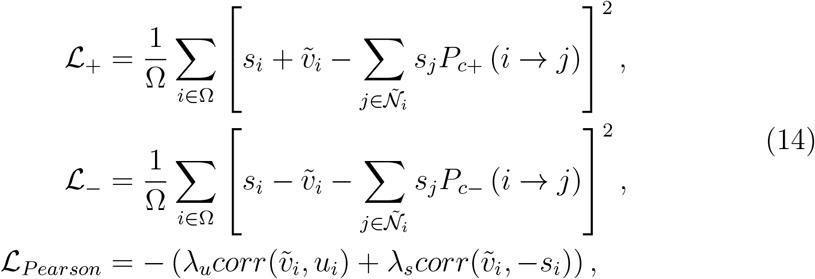

where *corr* denotes the Pearson correlation coefficient. We use the combination of the objective terms 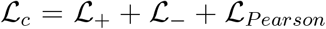 to train the DeepVelo model. *λ_u_, λ_s_* are constant factors to balance the scale of objective terms. The model parameters are optimized to minimize the 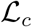.

Notably, for each gene, the optimization **integrate the information of other genes**, because the target cell probability estimation of *P*(*i → j*) considers the full gene expression of cell *i* and *j*. From a per gene estimate perspective, it corrects the target cell probability when the unspliced/spliced counts of the current gene are noisy, but the majority of genes point to the correct target cell *j*. This integration of genes is a unique advantage of DeepVelo compared to existing methods, and it particularly contributes to the capability of celltype-specific velocity prediction and time-dependent gene correction of DeepVelo (Section 3.3).

The DeepVelo model is trained by gradient back-propagation using the Adam [14] optimizer up to 100 epochs. The updated model at the last epoch is used to compute the estimated velocities.

### 5.4 Overall and celltype-wise consistency evaluation

The overall consistency score is proposed as the average cosine similarity of the velocity vectors to their neighbors. For each cell *i*,

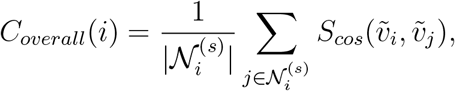

where 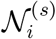 is the 30-nearest-neighbor cells with similar spliced gene expression, computed in the preprocessing step (Online Methods – 5.1). *S_cos_* denotes the cosine similarity operation. *v_i_,v_j_* are the estimated velocities from Eq.4.

The celltype-wise consistency computes the similarities within each celltypes instead. For each cell *i* and the celltype 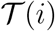 it belongs to,

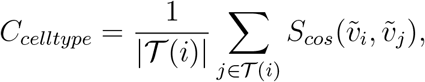

where 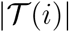 denotes the number of cells belonging to the celltype.

### 5.5 Computing cell-to-cell connectivity graph

The similarity of velocity vectors of cells could model cell-to-cell connectivities. We use the connectivity graph for downstream tasks, including driver gene analysis and developmental trajectory inference.

The weight in the connectivity graph, *w_ij_* denotes the estimated magnitude of connection. Higher *w_ij_* means the future state of cell *i* is close to the current state of cell *j*. *w_ij_* could be computed by possible similarity measures between velocity *v_i_* and the gene expression difference *s_j_* — *s_i_*. Here, we used the cosine similarity, which is also adopted in scVelo [4], therefore,

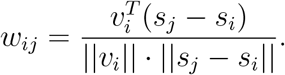

For the visualization of the velocity plot, we adopted the same projection computation provided by exiting methods [16, 4] to project velocity as arrows onto low-dimensional embeddings, such as tsne [28] and UMAP [21]. To summarize, the transition probability *π_i,j_* from a cell *i* to possible target cell *j* is computed by the Gaussian normalized connectivity weight *w_i,j_*. Then the velocity vector for *v_i_* in a low-dimensional space is computed by the weighted sum of 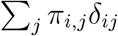, where *δ_ij_* is the direction vector pointing from cell *i* to *j* in the low-dimensional space.

### 5.6 Driver gene estimation and comparison

To determine functional signals in the driver genes, the top 100 genes based on a correlation with each lineage were determined, in particular for the hindbrain developmental data from [30]. Overlap with marker genes based on the original analysis used to annotate celltypes was performed, as well as overlap with transcription factors. Transcription factors were pulled from the manually annotated Human Transcription Factors list curated by Lambert et al. [17], and were lifted over to mouse data using orthologous gene-matches.

Analysis of marker overlap was further extended by determining the ranking of marker genes across all tested driver genes (2000 total) for both scVelo and DeepVelo per lineage in the Vladoiu et al. [30] data. The DeepVelo and scVelo predicted rankings of these marker genes for both lineages were compared, where a higher ranking of marker genes indicated a stronger signal for biologically relevant genes in the driver gene analysis. Since the entire tested driver gene lists were used, the number of genes per lineage was equivalent, and the rankings of the two lists were compared using the Mann-Whitney U Test (or Wilcoxon Rank-Sum Test), which is a non-parametric test for differences in sample distributions. The two-sided version of the test was used in this case, allowing either DeepVelo or scVelo to have greater or lesser rankings for relevant marker genes.

### 5.7 Pathway enrichment analysis

To determine functional signals in the driver gene results, pathway enrichment analysis was done using the ActivePathways R package [22]. The top 100 driver genes, based on correlation values for both the GABAergic and gliogenic lineages from the Vladoiu et al. [30] data, were input into the ActivePathways geneset enrichment analysis model. The latest Gene-Matrix-Transposed (GMT) files containing gene-set information from the Gene Ontology Molecular Function, GO Biological Process, and REACTOME databases were used [7, 13]. Pathways were labelled as being involved in “Neurogenesis”, ‘‘Developmental non-neuronal”, and “Non-specific” using manual annotation and the presence of known terms (such as “neuron projection” or ‘proliferation” for “Neurogenesis” and ‘Developmental non-neuronal”, respectively). “Non-specific” pathways indicated those that did not have immediately obvious roles in either neurogenesis or general development. To determine significant differences between pathway labelling and potential enrichment of neurogenic/development specific pathways, a two-sided Fisher’s exact test based on the hypergeometric distribution was done for the contingency table comprising of scVelo and DeepVelo pathway results and functional labels (“Neurogenesis”, ‘Developmental non-neuronal”, “Non-specific”) for the gliogenic and GABAergic lineages independently.

## 6 Acknowledgements

We thank Lin Zhang, Mehran Karimzadeh, Phil Fradkin, and Shihao (Rex) Ma of the Bo Wang Lab, and Jiao Zhang and Liam Hendrikse of the Michael Taylor Lab for their insightful comments on the manuscript.

## I Supplementary

**Figure S1:**
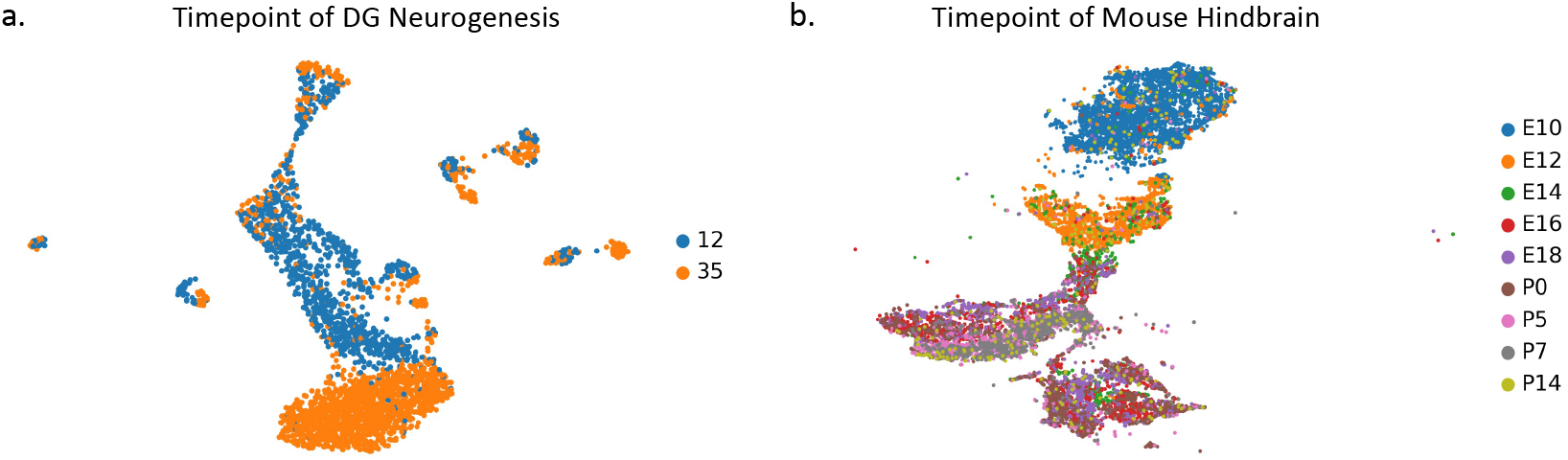
The developmental timepoints of sequenced cells in dendate gyrus neurogenesis (a) and mouse hindbrain development (b) datasets.

**Figure S2:**
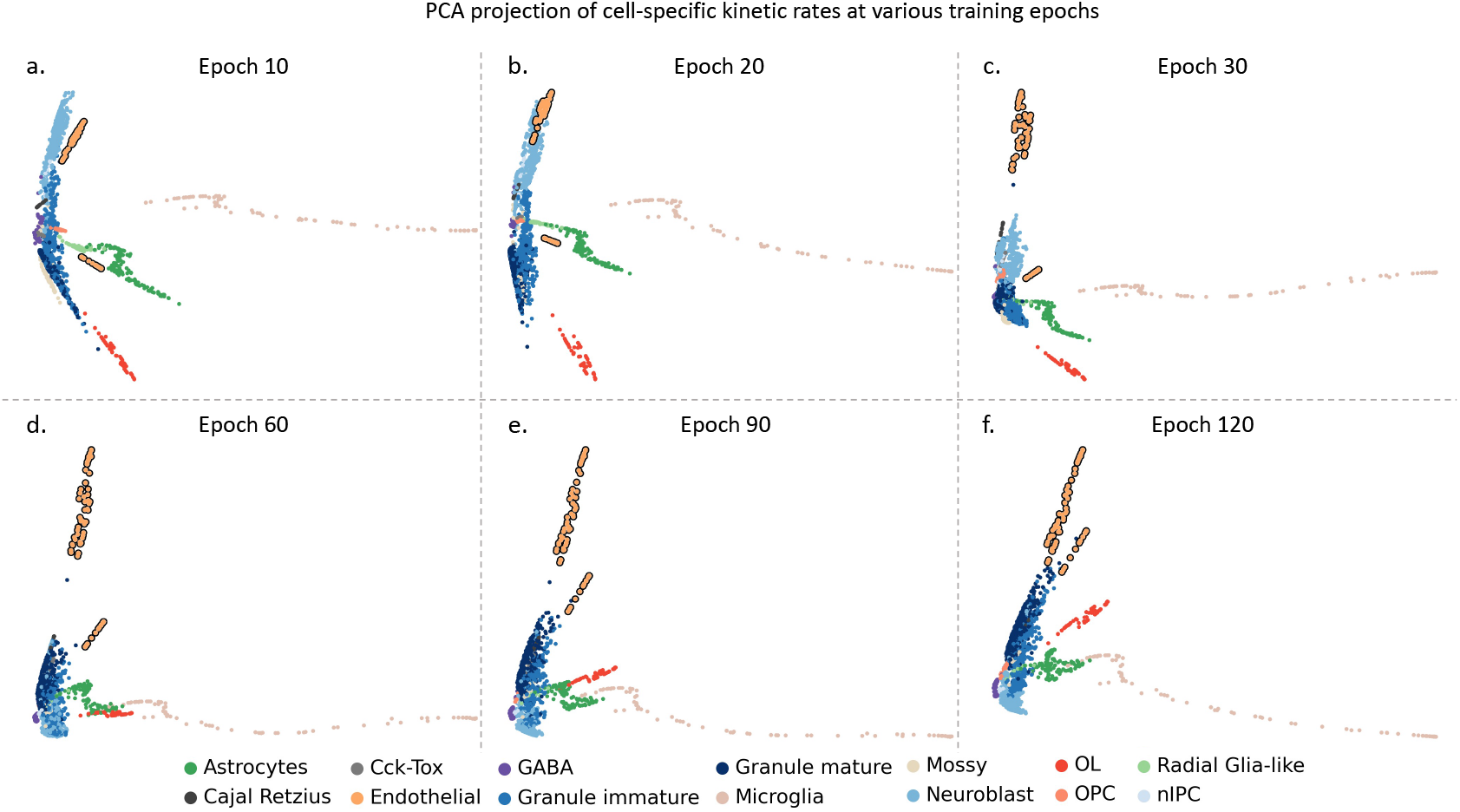
The PCA projection of cell-specific kinetic rates at various training epochs. (a-f) Scatter plot of the first two PCA dimensions at training epochs 10, 20, 30, 60, 90, 120. DeepVelo learns to predict similar kinetic rates for cells of same celltype. For example, the kinetic rates of Endothelial cells (outlined) are gradually clustered together and are located away from the unrelated granule lineage.

**Figure S3:**
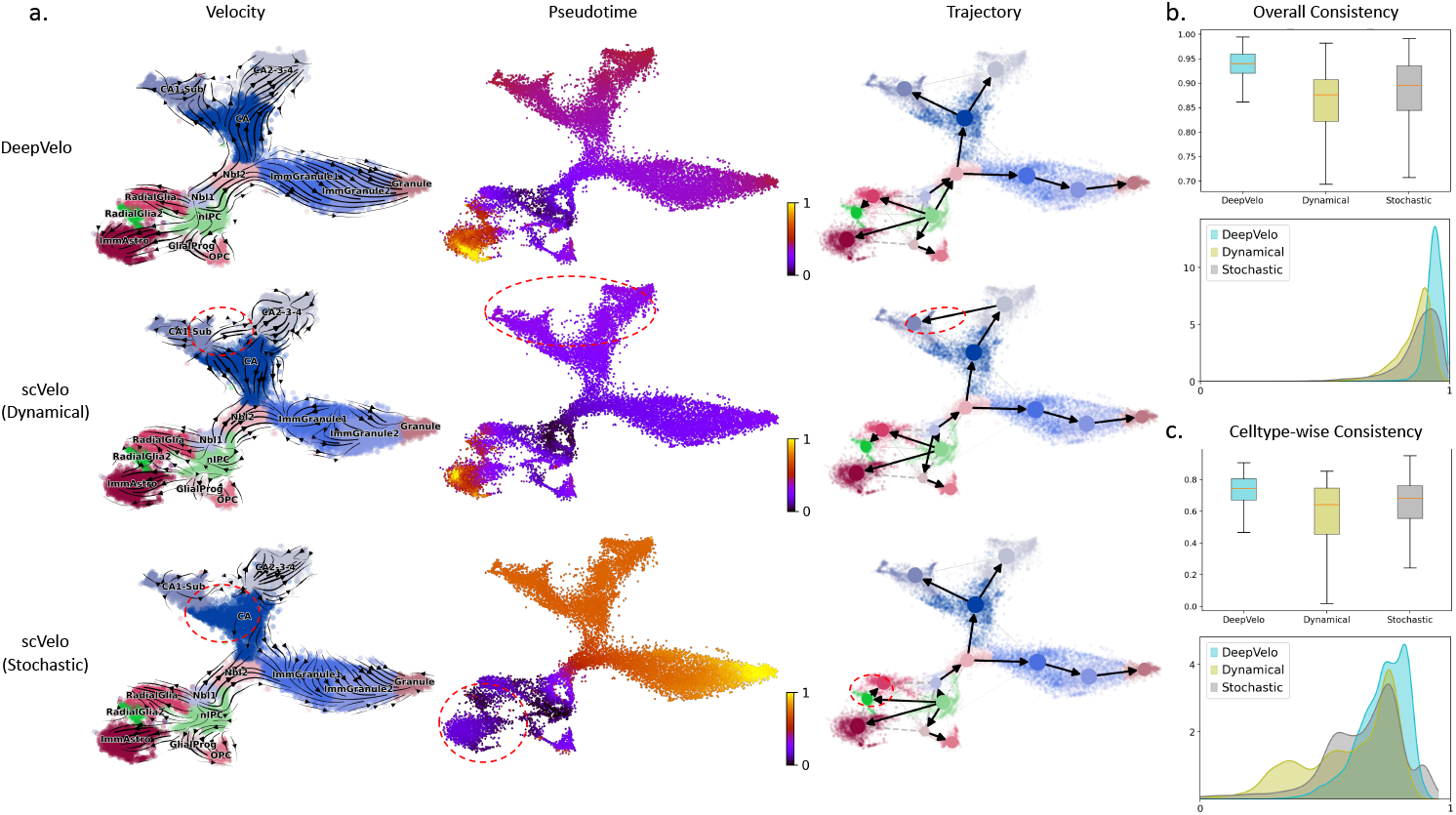
Comparison of three velocity methods on large-scale dentate gyrus data. (a) The velocity plot, pseudotime and trajectory inference of DeepVelo, scVelo dynamical model and scVelo stochastic mode, respectively. We highlighted observable incorrect predictions of compared methods in red circles. (b, c) The overall consistency score and celltype-wise consistency score. DeepVelo shows better performances regarding both metrics.

**Figure S4:**
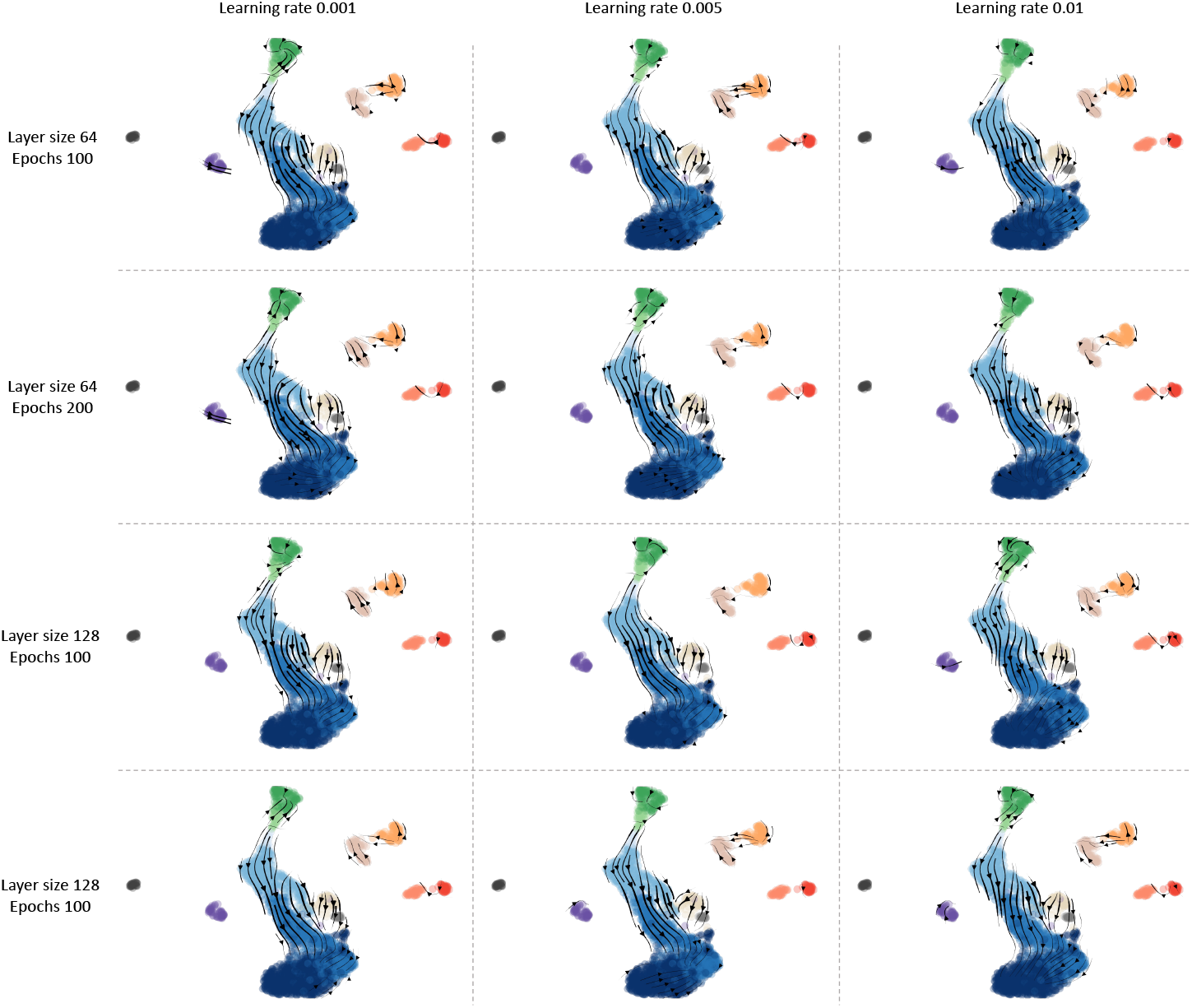
DeepVelo’s robustness with respect to key hyperparameters. Using different combinations of important hyperparameters, the DeepVelo velocity plots on the dentate gyrus neurogenesis data are depicted. DeepVelo consistently captures the correct velocity directions with respect to different learning rates, GCN layer size and number of training epochs.

**Figure S5:**
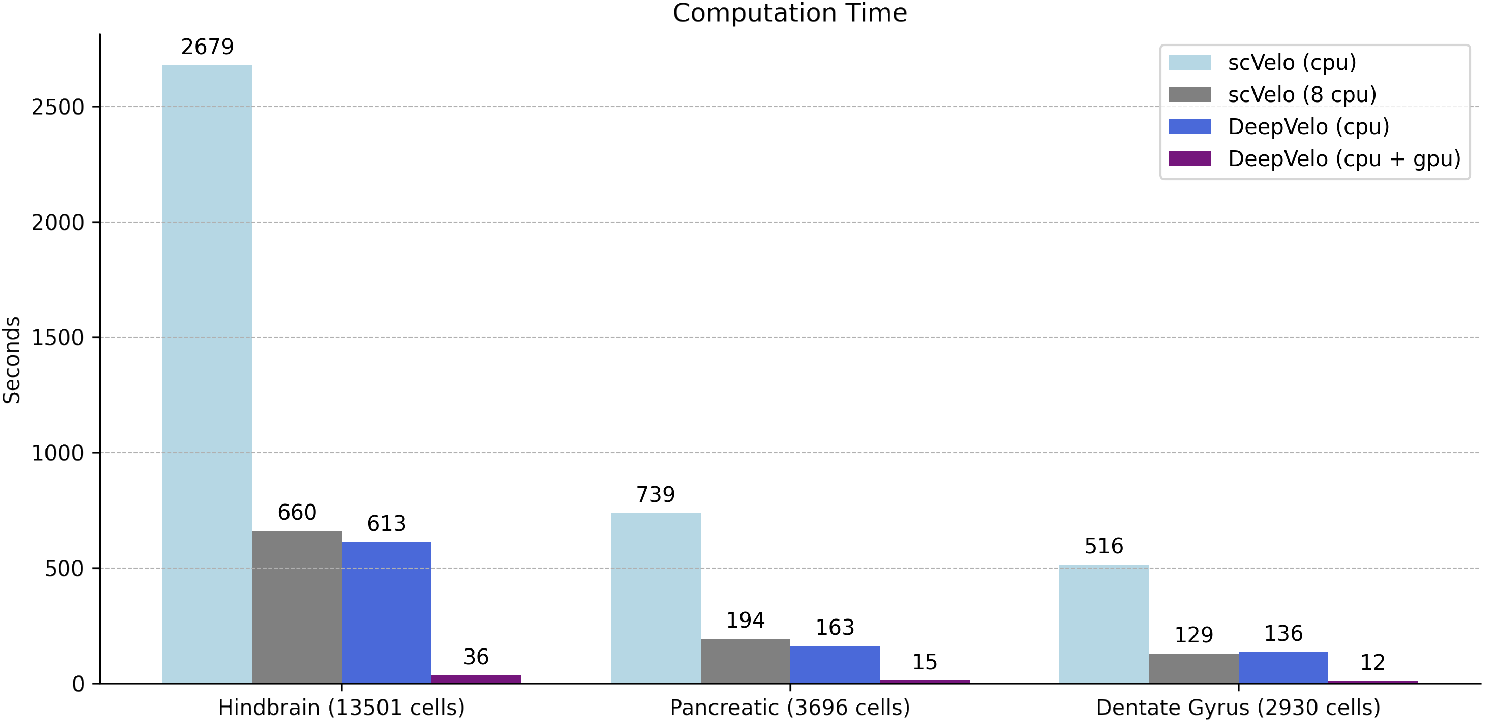
Computational efficiency comparison of scVelo and DeepVelo across datasets. Using the same CPU device(*), DeepVelo had a 4 fold acceleration compared to the dynamical model. Using GPUs, DeepVelo can complete training and estimation for over 13,000 cells in 36 seconds. Generally the GPU-accelerated DeepVelo is 10-20 times faster than the accelerated dynamical model (8 CPUs). (*) The DeepVelo(CPU) uses the pytorch package, which automatically utilizes 8 CPUs for the gradient optimization step. For all other computations, the DeepVelo(CPU) runs on single CPU.

**Figure S6:**
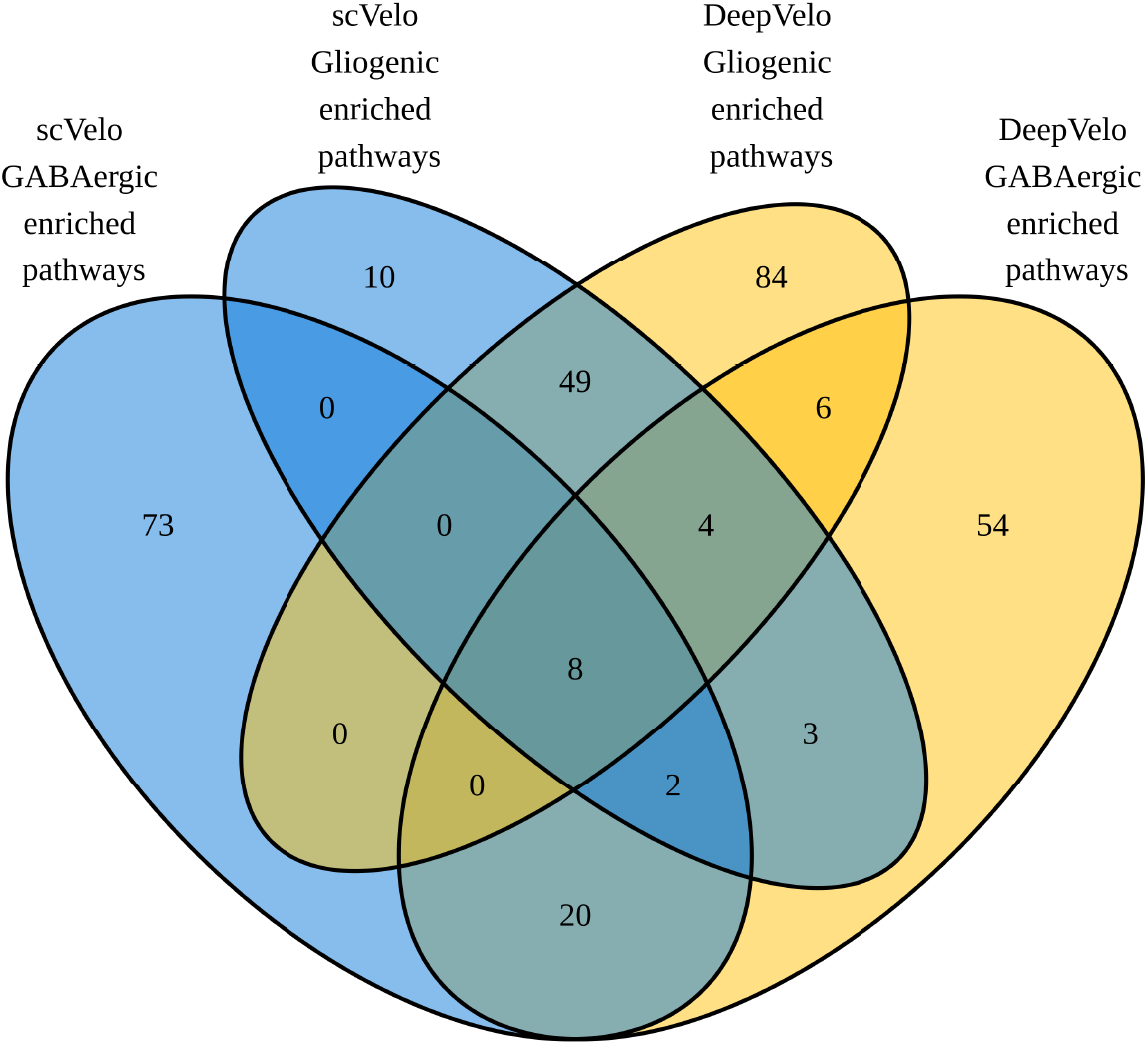
Full pathway enrichment analysis results overlap. Overlap of scVelo and DeepVelo pathway enrichment analysis results, between methods, for the top 100 GABAergic and gliogenic driver genes.

## Notes

https://github.com/bowang-lab/DeepVelo

### Competing Interest Statement

The authors have declared no competing interest.

### Summary of Updates

Results are revised using the updated method described in Online Methods

